# Advancing Thalamic Nuclei Segmentation: The Impact of Compressed Sensing and FastSurfer on MRI Processing

**DOI:** 10.1101/2024.07.05.602237

**Authors:** Sebastian Hübner, Stefano Tambalo, Lisa Novello, Tom Hilbert, Tobias Kober, Jorge Jovicich

## Abstract

The thalamus is a collection of gray matter nuclei that play a crucial role in sensorimotor processing and modulation of cortical activity. Characterizing thalamic nuclei non-invasively with structural MRI is particularly relevant for patient populations with Parkinson’s disease, epilepsy, dementia, and schizophrenia. However, severe head motion in these populations poses a significant challenge for in vivo mapping of thalamic nuclei. Recent advancements have leveraged the compressed sensing (CS) framework to accelerate structural MRI acquisition times in MPRAGE sequence variants, while fast segmentation tools like FastSurfer have reduced processing times in neuroimaging research.

In this study, we evaluated thalamic nuclei segmentations derived from six different MPRAGE variants with varying degrees of CS acceleration (from about 9 to about 1 minute acquisitions), using both FreeSurfer and FastSurfer for segmentation. Our findings show minimal sequence effects with no systematic bias, and low volume variability across sequences for the whole thalamus and major thalamic nuclei. Notably, CS-accelerated sequences produced less variable volumes compared to non-CS sequences. Additionally, segmentations of thalamic nuclei by FreeSurfer and FastSurfer were highly comparable.

We provide first evidence supporting that a good segmentation quality of thalamic nuclei with compressed sensing T1-weighted image acceleration in a clinical 3T MRI system is possible. Our findings encourage future applications of fast T1-weighted MRI to study deep gray matter. CS-accelerated sequences and rapid segmentation methods are promising tools for future studies aiming to characterize thalamic nuclei in vivo at 3T in both healthy individuals and clinical populations.

## 1. Introduction

The thalamus is a diencephalic structure of the mammalian forebrain involved in gating sensorimotor input to the cortex and modulating cortical activity via transthalamic cortico-cortical pathways (Jones, 2007; Sherman, 2017). It is composed of a collection of grey matter nuclei, each of which is characterized by specific histological (Jones, 2007; Morel, 2007; Sherman, 2017), connectional (Behrens et al., 2003; Sherman, 2017), and functional properties (Herrero et al., 2002; Zhang et al., 2008). Individual thalamic nuclei are differently associated with sensorimotor functions (Jones, 2007; Sherman, 2017) and cognitive processes, such as attention (Grieve et al., 2000; Guedj & Vuilleumier, 2023), memory (Leszczyński & Staudigl, 2016; Sweeney-Reed et al., 2021), emotions (Arend et al., 2015; Golden et al., 2016), language (Wahl et al., 2008; Hebb & Ojemann, 2013), executive functions (Van der Werf et al., 2003; Jakab et al., 2012), and consciousness (Schiff, 2008; Ward, 2011), and are also affected in neurological and psychiatric conditions including, among others, Alzheimer’s disease (de Jong et al., 2008; Power& Loi, 2015), Parkinson’s disease (Blesa et al., 2016; Wang et al., 2022), different forms of epilepsy (Natsume et al., 2003; Vetkas et al., 2022), schizophrenia (Buchsbaum et al., 1996; Byne et al., 2009), and obsessive-compulsive disorder (Van den Heuvel et al., 2016; Weeland et al., 2022).

Magnetic resonance imaging (MRI)-based structural and volumetric characterization of thalamic nuclei in humans has become increasingly important for both basic research and clinical purposes (Lozano, 2000; Iglesias et al., 2018; Keun et al., 2021). Nevertheless, *in vivo* mapping of thalamic nuclei can present technical challenges, since thalamic nuclear boundaries are notoriously difficult to visualize, for instance, even in standard T1-weighted (T1w) MRI (Magnotta et al., 2000; Iglesias et al., 2018; Najdenovska et al., 2019; Su et al., 2019; Rushmore et al., 2022). Another challenge can result from structural and volumetric biases induced by high levels of head motion during MRI (Reuter et al., 2015; Baum et al., 2018; Zacà et al., 2018), such as those occurring in patients with tremor, Alzheimer’s diseases, and also healthy elderly adults (Van Dijk et al., 2012; Iglesias et al., 2017). While reducing scanning times can mitigate the probability of motion artifacts, it is also important to consider that undersampled techniques might be more sensitive to motion due to less data being collected. Therefore, in addition to reducing scan times to improve patient comfort, workflow, and throughput, adopting tools that allow for accurate segmentation of thalamic nuclei, even in the presence of small motion artifacts, can help address these challenges.

Compressed sensing (CS; Donoho, 2006) is one among various acceleration techniques recently proposed for MRI. By only sampling a subset of the *k*-space rather than its full grid, CS allows for a faster acquisition of high-resolution MRI data (Lustig et al., 2007; Pauly, 2008). CS has recently been applied to several T1w sequences, such as the standard vendor-provided magnetization-prepared rapid gradient echo (MPRAGE; Mugler & Brookeman, 1990) and its variant MP2RAGE (Marques et al., 2010). In general, reports are of comparable quality and volumetry to parent non CS- accelerated sequences in a number of brain structures (Mussard et al., 2018; Mair et al., 2019, 2020; Mönch et al., 2020; Dieckmeyer et al., 2021; Ferraro et al., 2022), with biases at higher accelerations. So far, the effect of CS acceleration of various MPRAGE sequence variants has not yet been investigated on the volumetry of thalamic nuclei specifically. The acquisition of CS-accelerated structural images has the potential to help the structural and volumetric characterization of thalamic nuclei in patients with high level of head motion, especially since research protocols often combine multimodal MRI techniques that require long runtime scans.

The accurate segmentation of thalamic nuclei is another important factor for basic and clinical research. Together with faster scans, a reduction of data processing times without sensible effects on the outcome variables becomes an appreciated factor especially when *in vivo* research employs large samples. Several brain segmentation tools have been proposed, among which FreeSurfer (Fischl et al., 2002; Fischl, 2012) is well-known and commonly used (Despotović et al., 2015). An alternative to FreeSurfer has recently been proposed, called FastSurfer (Henschel et al., 2020). FastSurfer is a deep learning-based and extensively-validated whole-brain segmentation tool able to replicate FreeSurfer analyses in about 1 hour, as opposed to FreeSurfer, which takes several hours. There is evidence indicating it as a robust alternative to FreeSurfer in cortical and subcortical structures (Henschel et al., 2020; Bloch & Friedrich, 2021; Kemenczky et al., 2022; Müller et al., 2023) including the thalamus (Opfer et al., 2023). However, it remains unknown how the performance of FastSurfer for thalamic nuclei segmentation may be affected by T1w acceleration strategies.

In this study, we aimed to evaluate the robustness of thalamic nuclei volumetry by examining the effects of two key variables: the use of different MPRAGE acquisition variants with varying CS acceleration factors, and the application of thalamic nuclei segmentation methods with different processing speeds. Specifically, we compared thalamic nuclei segmentations obtained using FreeSurfer and FastSurfer across several MPRAGE variants, including standard MPRAGE (5:32 min), multi-echo MPRAGE (6:03 min), MP2RAGE (8:52 min), CS-MP2RAGE (3:40 min), and two CS-

MPRAGE variants (2:04 min and 1:14 min).

## 2. Materials and Methods

### 2.1 Participants

In the present study, GeneRalized Autocalibrating Partial Parallel Acquisition (GRAPPA)-accelerated sequences (Griswold et al., 2002) are referred to as non CS sequences. We used data from two samples: i) Sample A with both non CS and CS sequences: 15 healthy adults (mean age (SD) = 25.7 (3.5), range = [20.7, 32.5] years; 47% males), and ii) Sample B with only non CS sequences: 7 healthy adults (mean age (SD) = 30.2 (6.5), range = [24.1, 38.6] years; 57% males). No statistically significant differences in age and gender of the two samples was detected with non-parametric analyses.

### 2.2 MRI acquisition

Table 1 lists acquisition parameters of all the considered sequences. All subjects underwent 3T MRI (MAGNETOM Prisma, Siemens Healthcare, Erlangen, Germany) with a 64-channel head-neck radio-frequency receive coil. 3D T1w sequences were acquired with 1 mm isotropic voxels, same spatial coverage, prescan normalize on, and no image filters: MP2RAGE (TR / TE = 5000 / 2.98 ms; TI = 700 / 2500; α = 4° / 5°; GRAPPA = 3; TA = 8:52 min), multi-echo MPRAGE (meMPRAGE; TR = 2530 ms, TE_1-4_ = 1.69 / 3.55 / 5.41 / 7.27 ms, TI = 1100 ms, α = 7°; GRAPPA = 2; TA = 6:03 min), “standard” MPRAGE (TR / TE / TI = 2310 / 3.48 / 1200 ms; α = 12°; GRAPPA = 2; TA = 5:32 min; Mugler & Brookeman, 1990), research application CS-MP2RAGE (TR / TE = 5000 / 2.88 ms; α = 4° / 5°; samples/TR = 195; undersampling factor = 4.6; regularization factors = 0.0006 / 0.0004; TA = 3:40 min; Mussard et al., 2020), and two CS-MPRAGE (TR / TE / TI = 2300 / 2.88 / 900 ms; α = 9°; samples/TR = 196; regularization factor = 0.0006; Mussard et al., 2020) with different acceleration factors: i) undersampling factor = 3.6; TA = 2:04 min, and ii) undersampling factor = 6.6; TA = 1:14 min. More details on CS reconstructions can be found in Mussard et al. (2020). For the meMPRAGE image reconstruction, all echoes were averaged to yield one T1w image for the segmentations (Van der Kouwe et al., 2008). For the MP2RAGE reconstruction, a regularization was applied on the uniform image to remove typical noise in the background (O’Brien et al., 2014).

**Table 1:**
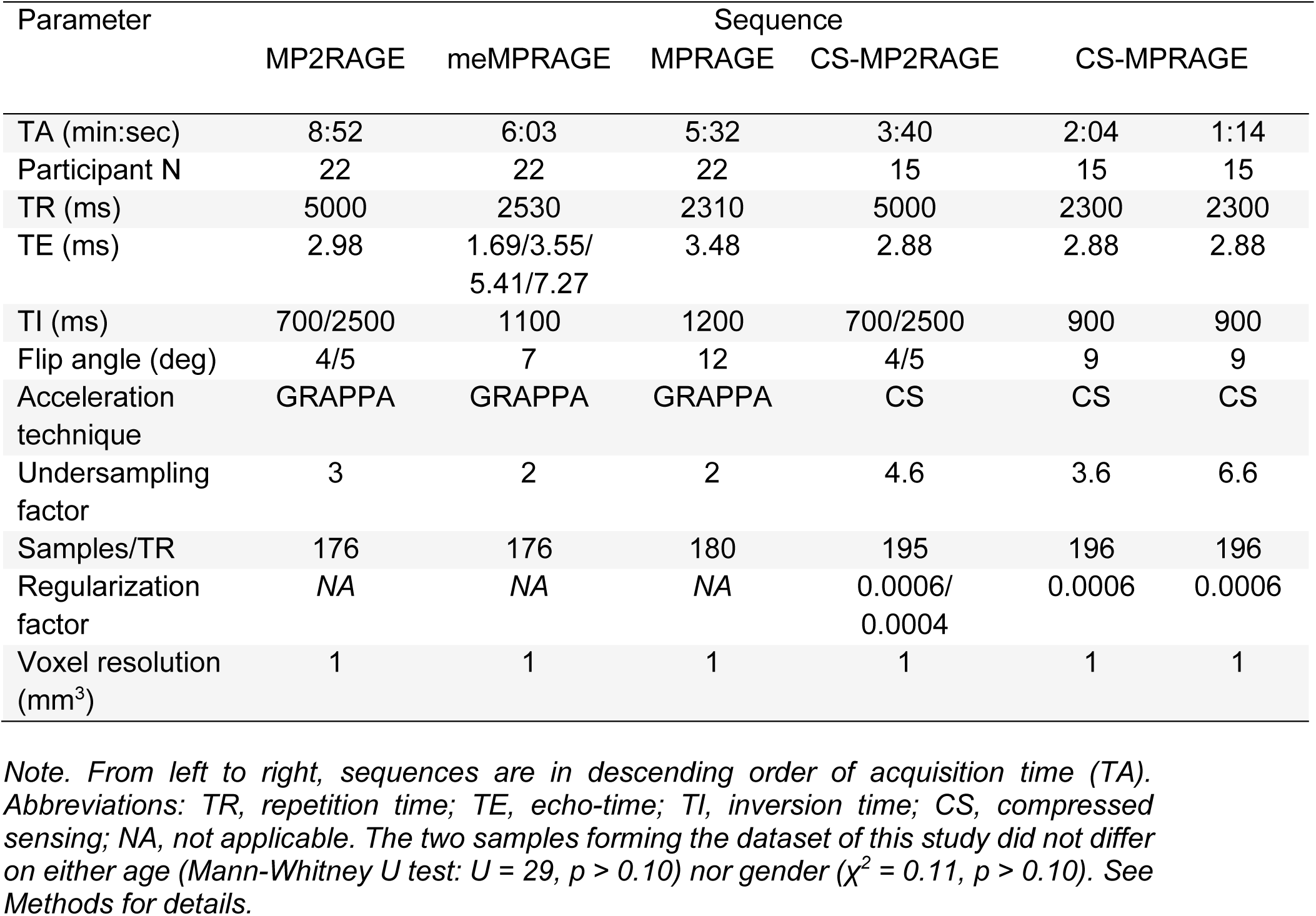
MRI parameters. Outline of MRI sequence parameters for the 3D MPRAGE variants used.

| Parameter | Sequence |  |  |  |  |  |
| --- | --- | --- | --- | --- | --- | --- |
|  | MP2RAGE | meMPRAGE | MPRAGE | CS-MP2RAGE | CS-MPRAGE |  |
| TA (min:sec) | 8:52 | 6:03 | 5:32 | 3:40 | 2:04 | 1:14 |
| Participant N | 22 | 22 | 22 | 15 | 15 | 15 |
| TR (ms) | 5000 | 2530 | 2310 | 5000 | 2300 | 2300 |
| TE (ms) | 2.98 | 1.69/3.55/<br>5.41/7.27 | 3.48 | 2.88 | 2.88 | 2.88 |
| TI (ms) | 700/2500 | 1100 | 1200 | 700/2500 | 900 | 900 |
| Flip angle (deg) | 4/5 | 7 | 12 | 4/5 | 9 | 9 |
| Acceleration technique | GRAPPA | GRAPPA | GRAPPA | CS | CS | CS |
| Undersampling factor | 3 | 2 | 2 | 4.6 | 3.6 | 6.6 |
| Samples/TR | 176 | 176 | 180 | 195 | 196 | 196 |
| Regularization factor | NA | NA | NA | 0.0006/<br>0.0004 | 0.0006 | 0.0006 |
| Voxel resolution (mm<sup>3</sup>) | 1 | 1 | 1 | 1 | 1 | 1 |

*Note. From left to right, sequences are in descending order of acquisition time (TA). Abbreviations: TR, repetition time; TE, echo-time; TI, inversion time; CS, compressed sensing; NA, not applicable. The two samples forming the dataset of this study did not differ on either age (Mann-Whitney U test: $U = 29$ , $p > 0.10$ ) nor gender ( $\chi^2 = 0.11$ , $p > 0.10$ ). See Methods for details.*

### 2.3 Whole-brain segmentation

T1w images were processed through the fully automated FreeSurfer (v. 7.1.1) *recon-all* stream, which performs all preprocessing steps, including motion and bias field corrections, as well as cortical and subcortical segmentation (Fischl et al., 2002; Fischl, 2012; https://surfer.nmr.mgh.harvard.edu).

Alongside FreeSurfer and for the purposes of quantitative volumetric comparison, we also employed the recent whole-brain segmentation tool FastSurfer (Henschel et al., 2020). The main distinctions between FreeSurfer and FastSurfer reflect their differences in computational steps: in contrast to FreeSurfer, which runs intensive computations like non-linear atlas registrations, brain extraction and intensity normalizations to achieve segmentations, FastSurfer utilizes convolutional neural networks (CNN) to recognize and segment cortical and subcortical structures, subsequently using them for cortical surface reconstruction, labeling, and thickness parametrization. More details about the rationale and workflow of FastSurfer can be found in the original publication and its online companion (Henschel et al., 2020; https://deep-mi.org/research/fastsurfer).

With the segmentation method FastSurfer, T1w images were segmented using the *FastSurferCNN* pipeline, which runs whole-brain volumetric segmentations and yields equivalent data to the *aparc.DKTatlas+aseg.mgz* file of FreeSurfer, as well as via the *recon-surf* stream, which creates surface-based cortical thickness data from the *FastSurferCNN* segmentation.

### 2.4 Thalamic nuclei parcellation

For both FreeSurfer and FastSurfer whole-brain segmentations, we further segmented the thalamus in its nuclear subdivisions using the thalamic parcellation tool developed by Iglesias et al. (2018; https://freesurfer.net/fswiki/ThalamicNuclei), which provides volumetric data for each thalamic nucleus.

In order to reduce the number of thalamic segmentation labels, we merged thalamic nuclei subdivisions as follows: pulvinar (PU; Subdivisions: PuM, PuA, PuL, PuI), ventrolateral (VL; Subdivisions: VLa, VLp), mediodorsal (MD; Subdivisions: MDm, MDl), ventral anterior (VA; Subdivisions: VA, VAmc); and intralaminar (IL; Nuclei: CM, CeM, CL, Pc, Pf). The following are further thalamic nuclei considered in the present study for which we have not changed labels with respect to Iglesias et al. (2018): anteroventral (AV), laterodorsal (LD), lateral geniculate nucleus (LGN), lateral posterior (LP), limitans-suprageniculate (L-Sg), medial geniculate nucleus (MGN), medial ventral-reuniens (MVRe), paratenial (Pt), and ventromedial (VM). We relabeled nucleus VPL to VP by virtue of name consistency with other major thalamic nuclei (Jones, 2007). Similar merging and relabeling schemes have been used by other authors as well (e.g., Iglesias et al., 2018; Bocchetta et al., 2020; Tregidgo et al., 2023). The merged volume of a given thalamic nucleus was computed as the sum of its subdivisions’ volumes, independently for each subject, sequence, segmentation tool, and hemisphere. For a complete description of nuclei colors and relabeling scheme see Supplementary Table 1.

### 2.5 Statistical analysis

Since we investigated within-subject volumetry across MPRAGE variants and did not compare individuals, we used raw thalamic nuclei volumes (mm^3^) without adjusting for intracranial volume or thalamus size. Within-subject volumetric variability across sequences was evaluated using coefficients of variation (CV), defined as the ratio of the standard deviation of volumetric data across sequences to its mean, then averaging CVs across subjects. CV computation was applied separately for the two segmentation tools considered in this study, and for each thalamic structure. We assessed the similarity of volume variability in left- and right-hemispheric data using the modified signed-likelihood ratio test for equality of CVs (Krishnamoorthy & Lee, 2014), in both FreeSurfer and FastSurfer data, in order to see whether the dimensionality of the study variables could be reduced by averaging volumes across hemispheres.

In order to explore the relation between thalamic nuclei size and volume variation, we computed Pearson’s correlation coefficient (*r*) between volumes and CVs. Furthermore, in order to evaluate the consistency in volume variability for the two considered segmentation tools, we calculated Pearson’s *r* between FreeSurfer and FastSurfer CVs in thalamic nuclei.

Friedman rank-sum tests were used to assess sequence effects on volumetric data of thalamic structures. Friedman test is the non-parametric equivalent of a repeated-measure one-way analysis of variance (Friedman, 1937). To investigate whether the proportion of sequence effects was different depending on the acceleration type (non CS *versus* CS accelerations), we used the paired samples McNemar’s test on proportions with Yates’ continuity correction for non CS and CS data. Furthermore, we assessed sequence effect size using Kendall’s W coefficients of agreement (Kendall, 1948; Tomczak & Tomczak, 2014). Kendall’s W is the normalized Friedman-test statistic and it ranges between 0 (no agreement) and 1 (complete agreement).

We also evaluated the agreement between FreeSurfer and FastSurfer thalamic nuclei volumes using Bland-Altman analysis (Bland & Altman, 1986), and the intraclass correlation coefficient (ICC, Fisher, 1936) as a measure of consistency between the two segmentation tools. For each thalamic structure, normalized mean differences were computed as the absolute value of the ratio of the difference between volumes of the two tools to their mean.

All statistical analyses were performed with software R (v 4.2.2; R Core Team, 2022).

## 3. Results

### 3.1 T1-weighted MRI Quality Assurance

Figure 1 shows T1-weighted images of a representative sample subject for qualitative comparison of MR images across MPRAGE variants. Image blurring increased expectedly with higher accelerations, though image contrast was well preserved across sequences.

**Figure 1:**
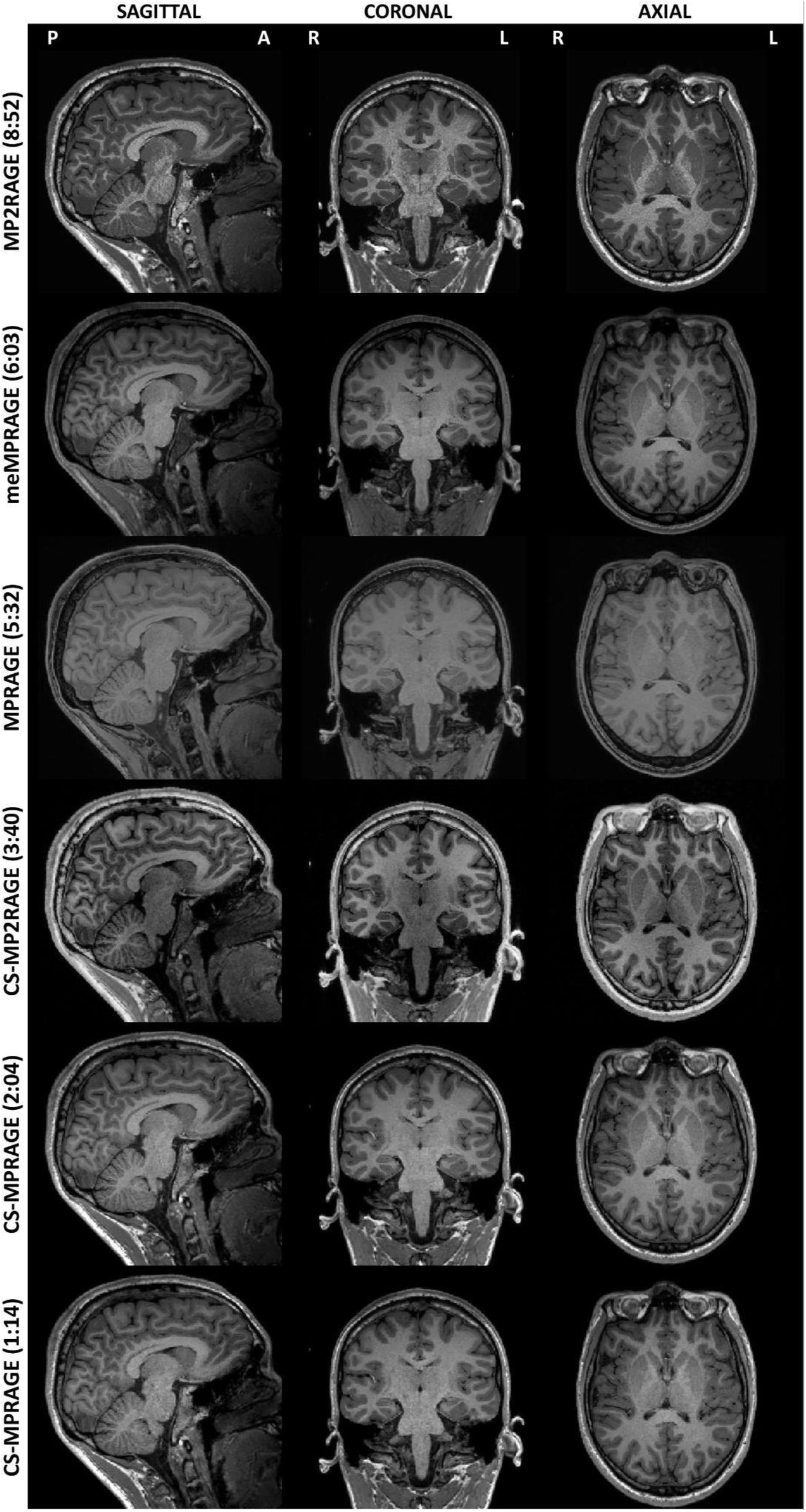
Structural T1w QA. T1-weighted image contrasts of differently accelerated MPRAGE variants. Qualitative comparisons of sagittal, coronal, and axial planes of a representative sample subject across the considered sequences (acquisition times in brackets, sequences details in Table 1). Image brightness parameters were adjusted to better show tissue contrasts. Slices are taken at central thalamus (subject’s native space). Sagittal images show the right hemisphere.

Figure 2 and Figure 3 show, respectively, a 3D rendering of thalamic nuclei and FreeSurfer automated thalamic parcellations across sequences in a representative sample subject. Visual inspection of the thalamic nuclei segmentations showed qualitative similarities for all MPRAGE variants, in segmentations provided by both FreeSurfer and FastSurfer tools. This confirmed that the range of accelerations evaluated have sufficient quality for enabling automated thalamic nuclei segmentations from both tools, FreeSurfer and FastSurfer. With this finding, we proceeded for a quantitative and systematic evaluation of how thalamic nuclei segmentation is affected by sequence and segmentation tools.

**Figure 2:**
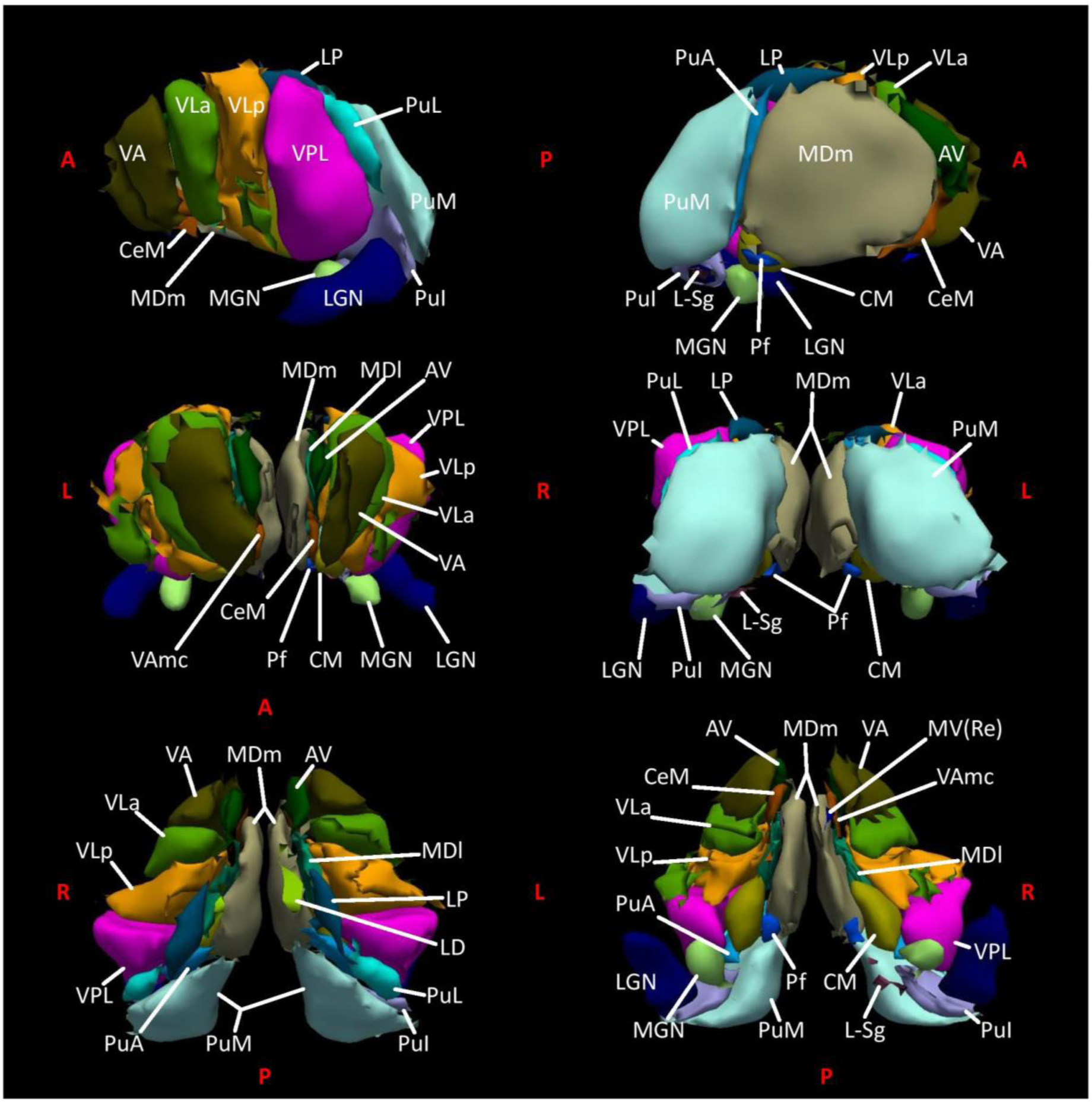
3D thalamic nuclei. Thalamic nuclei segmentation. FreeSurfer 3D rendering of thalamic nuclei in their lateral-medial (top row), rostral-caudal (middle row), and dorsal-ventral (bottom row) aspects on T1-weighted multi-echo MPRAGE data of a representative sample subject. In the present study, thalamic nuclei subdivisions were merged as follows: PU (PuM, PuA, PuL, PuI); VL (VLa, VLp); MD (MDm, MDl); VA (VA, VAmc); and IL (CM, CeM, CL, Pc, Pf). See Supplementary Table 1 for nuclei colors and details on the relabeling scheme.

**Figure 3:**
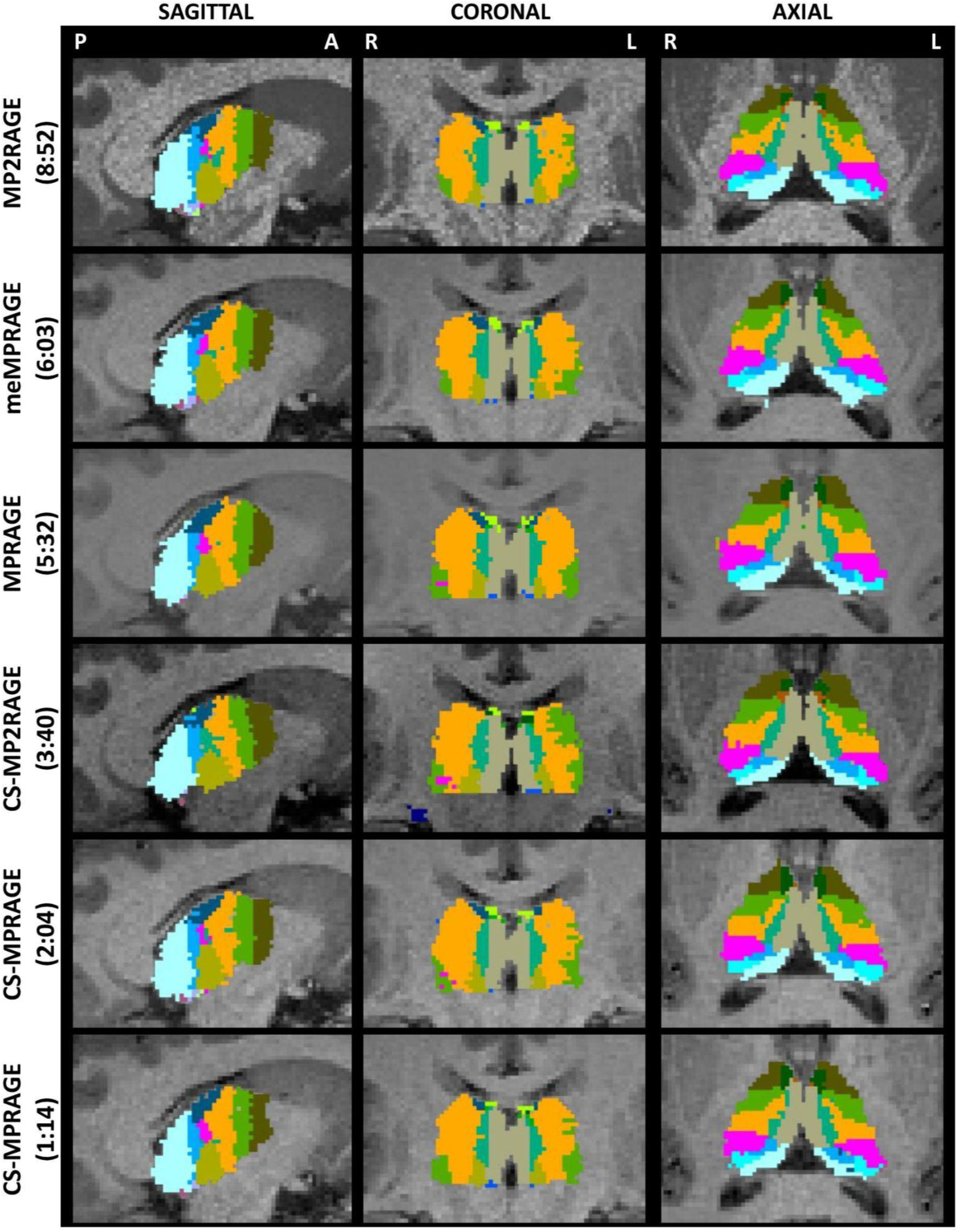
T1w thalamic nuclei segmentations QA. Thalamic nuclei segmentations for differently accelerated MPRAGE variants. FreeSurfer automated thalamic nuclei parcellations of a representative sample subject across the considered MPRAGE variants (acquisition times in brackets, sequences details in Table 1). Sagittal, coronal, and axial planes of parcellations are with the same brightness parameters as in Figure 1. As in Figure 1, slices are taken at central thalamus (subject’s native space). Sagittal images show the right hemisphere. Nuclei labels and colors as in Figure 2 and Supplementary Table 1. FastSurfer segmentations on the same subject’s native space can be seen in Supplementary Figure 1.

### 3.2 Within-Subject Sequence Effects

#### 3.2.1 Volume variability across sequences

We evaluated the level of within-subject volumetric variability across MPRAGE variants in the two segmentation tools FreeSurfer and FastSurfer. Table 2 lists volumes and coefficients of variation (CV) of thalamic nuclei across the various sequences (FreeSurfer data). Figure 4A and Figure 4B show, respectively, volumes and group-averaged within-subject CVs in thalamic nuclei, emphasizing how larger nuclei had similar, low variability (< 10%) regardless of acceleration, while smaller nuclei yielded lower variability with compressed sensing (CS) acceleration (Fig. 4C). Figure 4D shows that smaller nuclei were generally associated with higher volume variability across sequences, while Figure 4E shows that variability effects are consistent in both FreeSufer and FastSurfer data.

**Figure 4:**
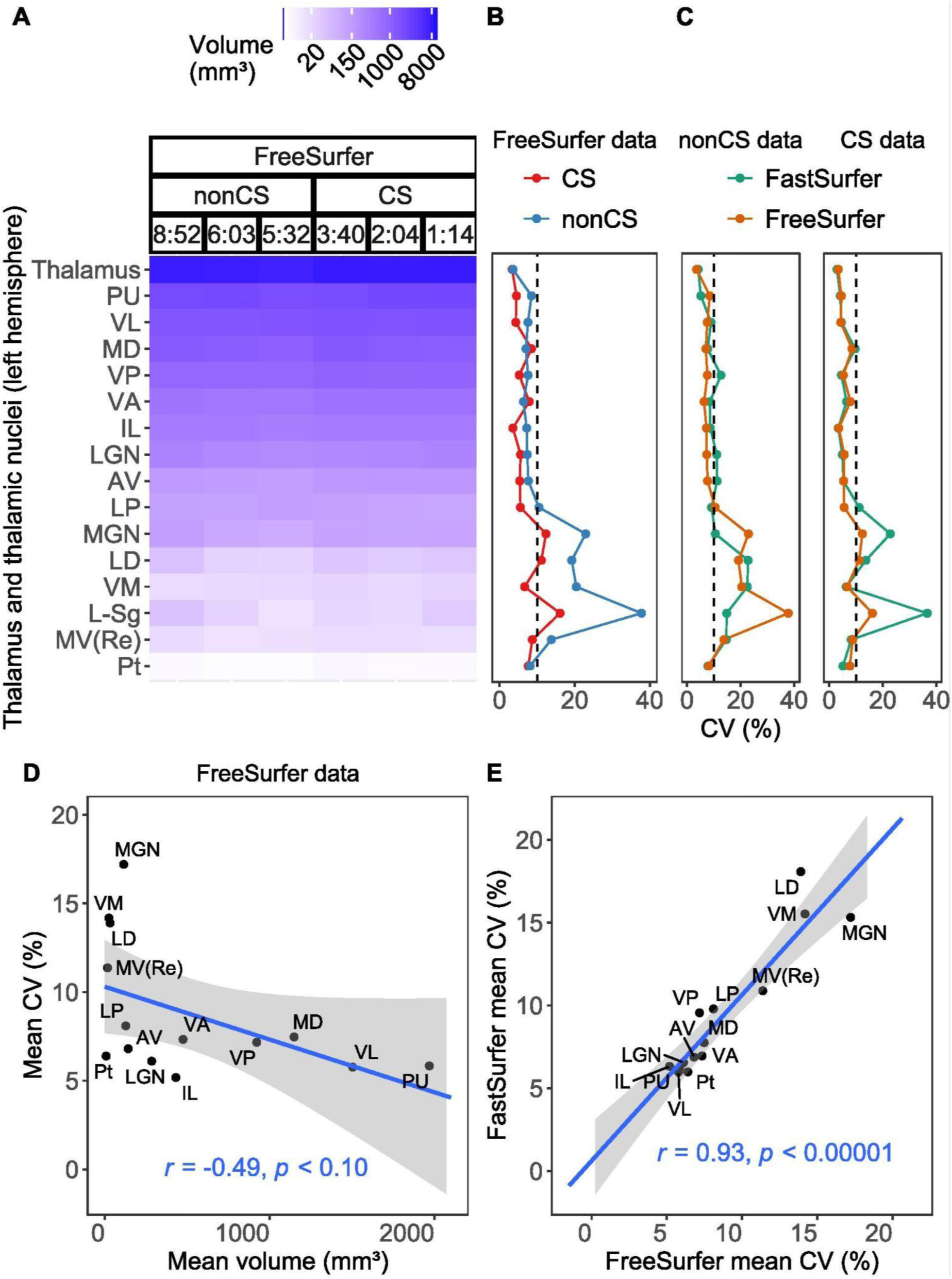
Volumes and CV. Within-subject volume variation in thalamic structures across MPRAGE variants. A) Thalamic volumes (logarithmic color-coding by size) segmented by FreeSurfer for each MPRAGE variant, grouped as non compressed sensing (nonCS) and compressed sensing (CS) accelerations, and listed by decreasing acquisition times (TA, minutes:seconds; 8:52, MP2RAGE; 6:03, meMPRAGE; 5:32, MPRAGE; 3:40, CS-MP2RAGE; 2:04 and 1:14, CS-MPRAGE). Left hemispheric data is presented, right- hemispheric data were similar; B) Within-subject coefficients of variation (CV) across MPRAGE sequences for thalamic structures (left and right hemispheres averaged), considering CVs from nonCS (blue) and CS (red) sequences separately. C) Within-subject CVs across sequences for thalamic structures (left and right hemispheres averaged), considering CVs from FreeSurfer (orange) and FastSurfer (green) segmentation tools separately, in nonCS (left hand-side) and CS (right hand-side) data. In panels B and C, vertical dash lines correspond to a reference CV value of 10%; D) Significant negative correlation between thalamic nuclei CVs and volume (data averaged across hemispheres, sequences, and segmentations). E) Significant positive correlation between CVs of FreeSurfer and FastSurfer segmentation tools (data averaged across hemispheres and sequences). In panels D and E, whole thalamus and L-Sg nucleus data were excluded (see text for details); including whole thalamus data in the linear fit did not alter results significantly for both associations (results not reported); Pearson’s r coefficients and their significance are presented; regression models show 95% confidence intervals of predictive values. Thalamic nuclei labels in Supplementary Table 1. FastSurfer volumes and volume variability in thalamic nuclei are shown in Supplementary Figure 2.

**Table 2:**
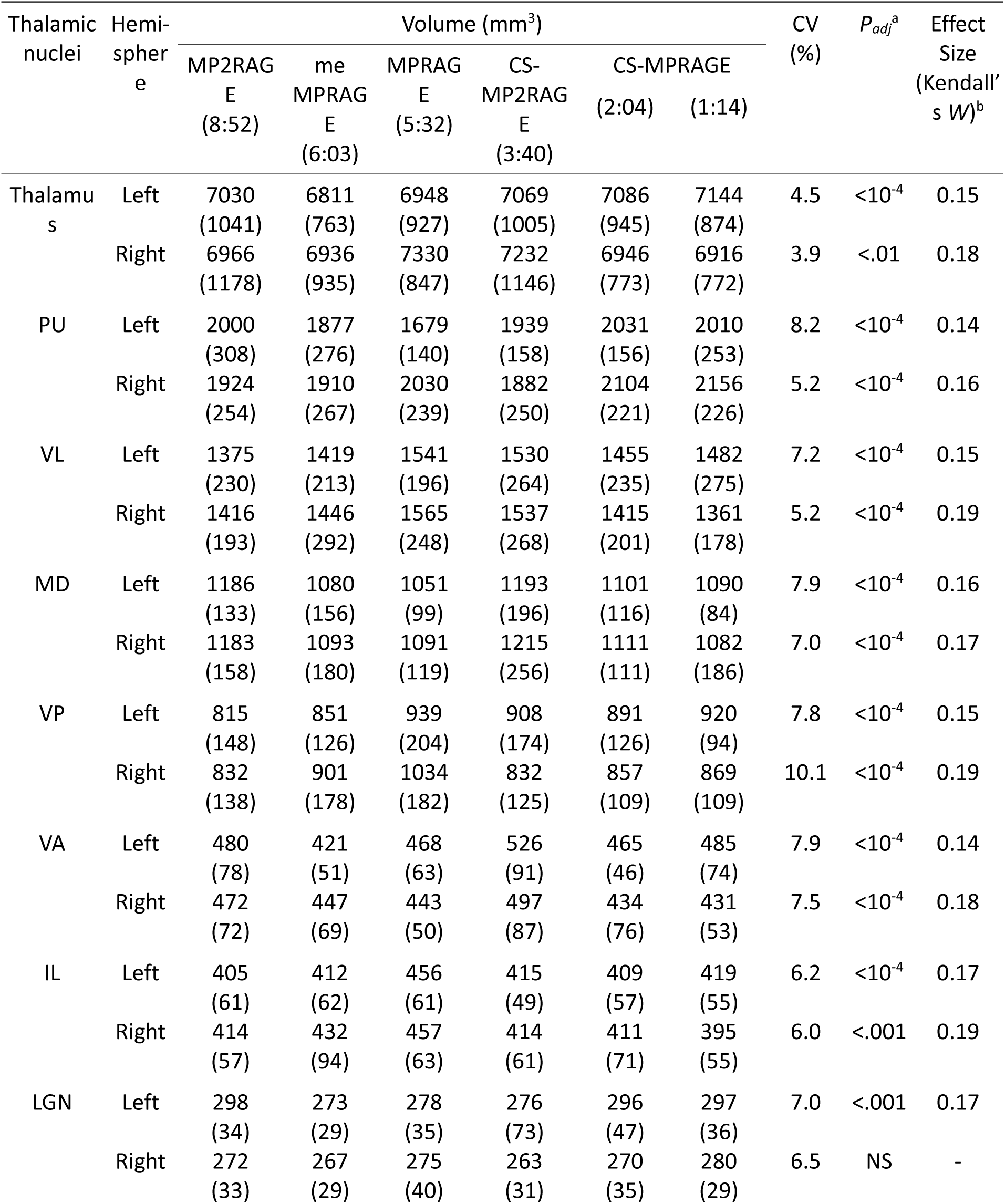
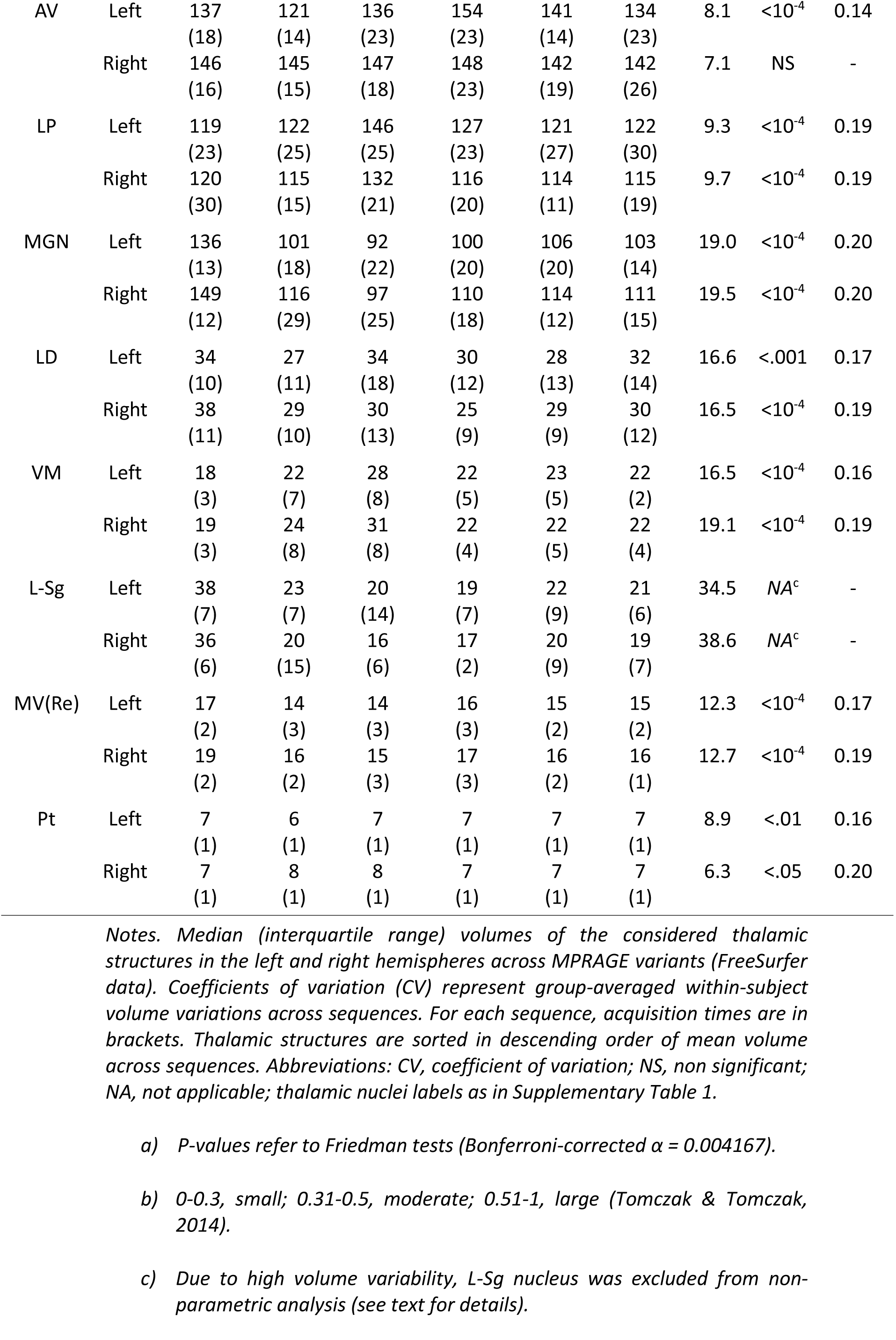
Thalamic volumes, CV, Kendall’s W. Volumes, volume variability, sequence effects, and sequence effect sizes of thalamic structures across MPRAGE variants. FreeSurfer volumetric segmentations for each MPRAGE sequence, group mean of within-subject coefficients of variation (CV) across MPRAGE variants, Friedman tests significance levels (adjusted p-values), and sequence effect sizes (Kendall’s W coefficients) for each thalamic structure and MPRAGE variant. Thalamic structures are sorted in descending order of mean volume across sequences.

##### 3.2.1.1 Whole thalamus

FreeSurfer data CVs in the whole thalamus were 4.5% and 3.9% in the left and right hemispheres, respectively, while 3.9% and 4.0% in FastSurfer data. We investigated potential hemispheric effects in thalamic volume variabilities for both FreeSurfer and FastSurferd data. The modified signed-likelihood ratio test for equality of CVs did not reveal significant hemispheric differences in either of the two considered segmentation tools. For this reason we averaged thalamic volumes across hemispheres.

##### 3.2.1.2 Thalamic nuclei

Considering FreeSurfer-data mean CVs across hemispheres, 10 out of 15 nuclei showed variability lower than 10%, four nuclei, namely MGN, LD, VM, and MV(Re), had variability in the range 10-20%, and the most variable nucleus across sequences was L-Sg (36.6%). Similarly, in FastSurfer data (Supplementary Figures 1 and 2), 8 out of 15 nuclei showed CVs lower than 10%, 6 out of 15 in the range 10-20%, and L- Sg was again the nucleus with the highest volume variation across sequences (22.9%). For this reason, L-Sg was excluded from further analyses.

As expected, we found volumetric variability across sequences to be inversely proportional to the volume of thalamic nuclei (Fig. 4D). Averaging across hemispheres and sequences, and excluding whole thalamus and L-Sg data, we found significant negative correlations between CVs and thalamic nuclei volumes in both FreeSurfer (Pearson’s *r*_(12)_ = -0.49, *t* = -1.97, *p* < 0.10, 95% CI [-0.81, -0.05]), and FastSurfer data (Pearson’s *r*_(12)_ = -0.47, *t* = -1.84, *p* < 0.10, 95% CI [-0.80, -0.08]).

Furthermore, we found a strong association between CVs of thalamic nuclei volumes from FreeSurfer and FastSurfer data (Fig. 4C). Excluding whole thalamus and L-Sg data, the linear fit was excellent (Pearson’s *r*_(12)_ = 0.93, *t* = 8.99, *p* < 0.00001, 95% CI [0.80, 0.98]), showing consistency in volume variability for these two segmentation tools.

#### 3.2.2 Sequence effects and effect sizes

Figure 5 shows whole thalamus and thalamic nuclei volumes derived from FreeSurfer for the considered MPRAGE variants. We investigated sequence effects in thalamic nuclei volumes across the various sequences, considering hemispheric data separately. Although Friedman tests yielded significant differences in nearly all thalamic nuclei, we observed no systematic bias in data according to sequence acceleration. In addition, a paired sample McNemar’s test on proportions with Yates’ continuity correction was performed to investigate marginal homogeneity of non CS and CS data. Results revealed that the proportion of sequence effects was significantly higher in non CS data as compared to CS (80% *versus* 33% of thalamic nuclei, respectively; *χ^2^*_(1)_ = 10.56, *p* < 0.01). Furthermore, we computed Kendall’s W coefficients as a measure of sequence effect size. All effect sizes were small (W range = [0.14, 0.20]).

**Figure 5:**
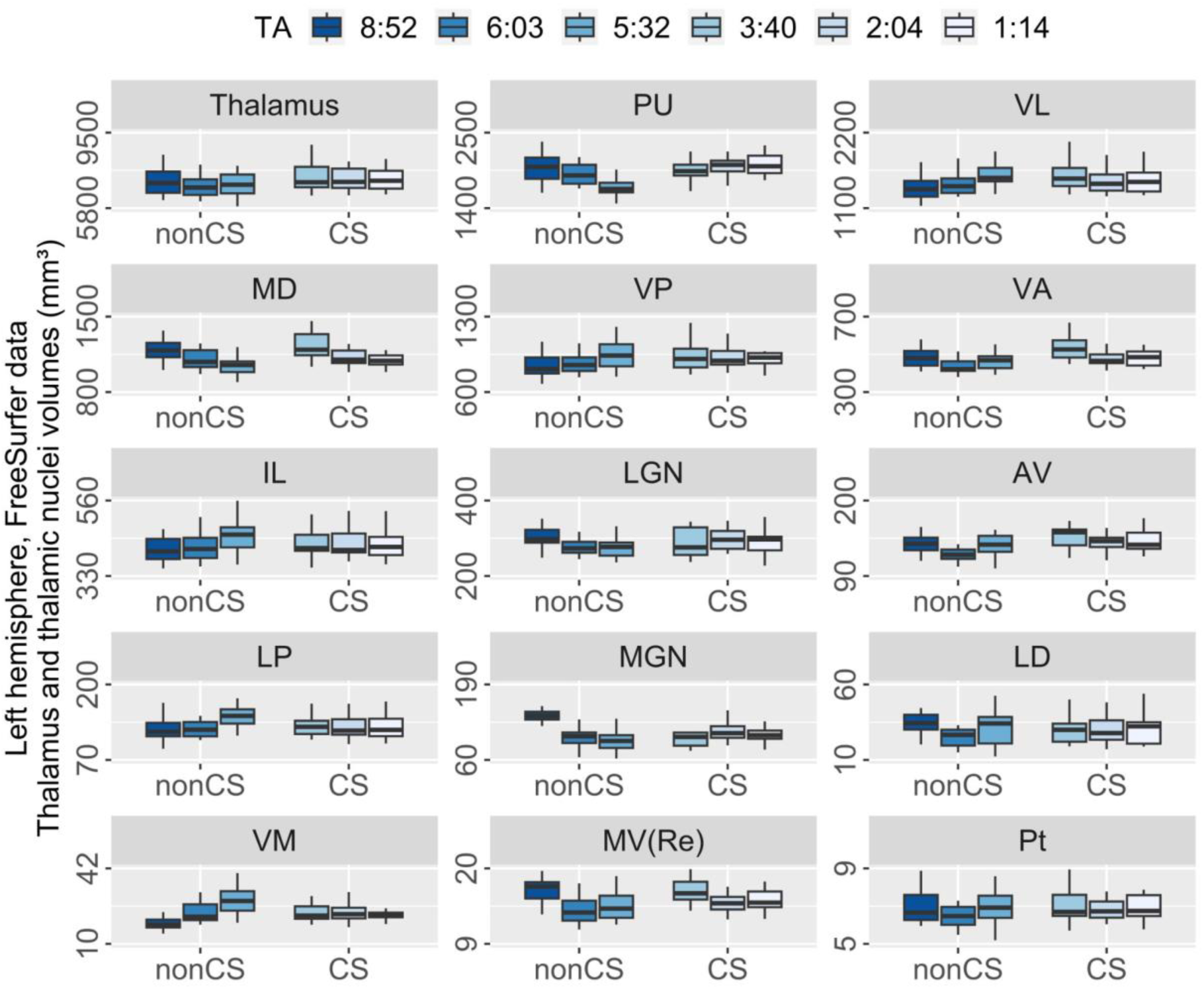
Boxplots of thalamic volumes in sequences. MPRAGE sequence effects on thalamic volumes. FreeSurfer volumetric segmentations in thalamic structures (group median and interquartile range, left hemisphere) for each MPRAGE sequence (see Table 1 for details on sequences). For each structure, box-and-whiskers are color-coded, sorted in descending order of acquisition time (TA), and grouped as non compressed sensing (nonCS) and compressed sensing (CS) accelerated sequences; Friedman tests yielded significant sequence effects for all thalamic structures in the left hemisphere (see Table 2). Right hemisphere was similar. Sequence TA (minutes:seconds): 8:52, MP2RAGE; 6:03, meMPRAGE; 5:32, MPRAGE; 3:40, CS-MP2RAGE; 2:04 and 1:14, CS-MPRAGE. Thalamic nuclei labels as in Supplementary Table 1.

Table 2 lists volumes, Friedman test results, and Kendall’s W coefficients in the considered thalamic structures and sequences.

### 3.3 Volume Correlations

To evaluate sequence biases in thalamic segmentations, we computed pairwise Pearson’s *r* coefficients for all MPRAGE sequence combinations (FreeSurfer data). All correlations were positive, ranging between 0.25 and 0.97. Figure 6 presents correlation matrices for pairs of sequences in the considered thalamic structures.

**Figure 6:**
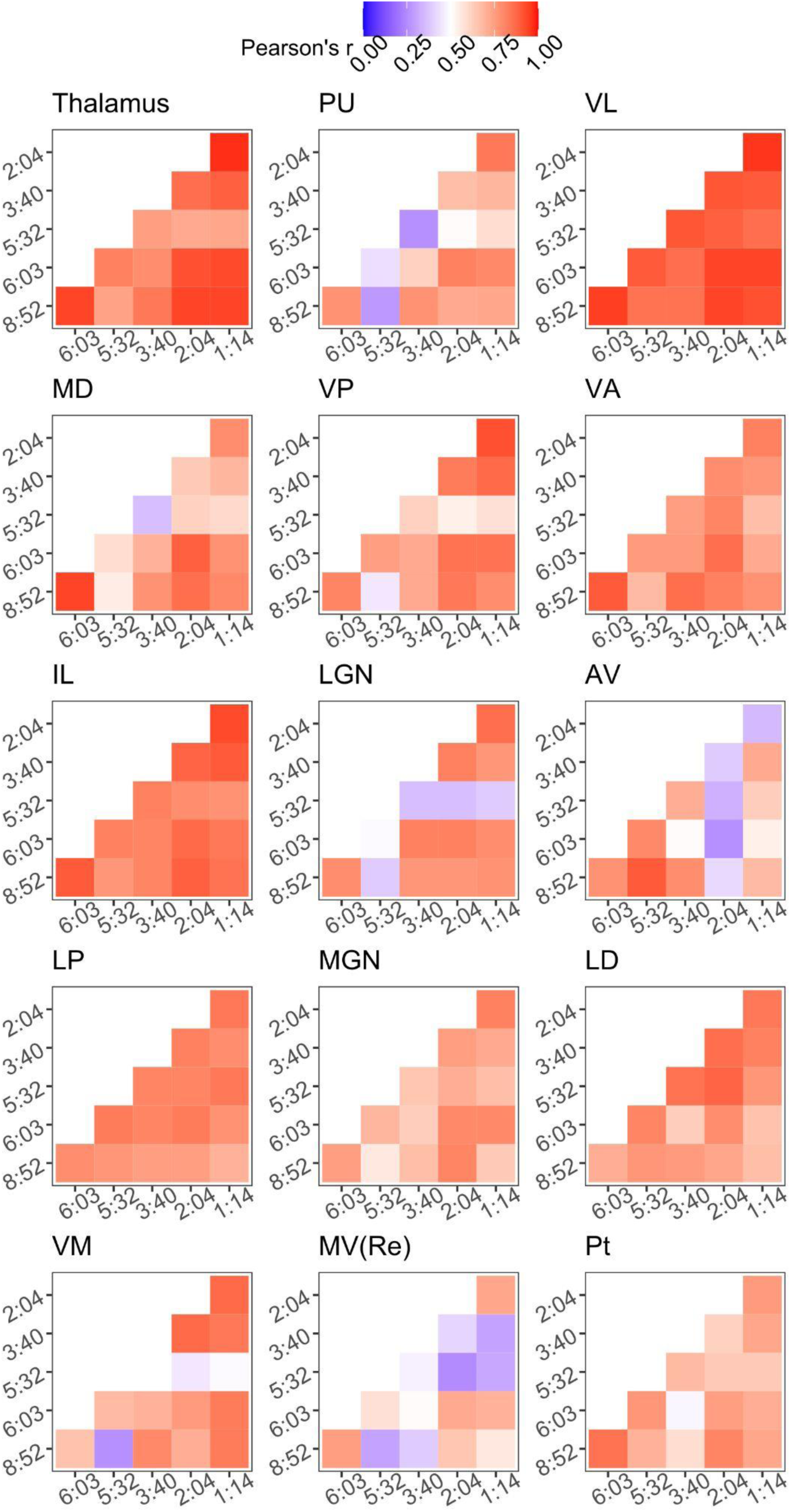
Pair-wise correlations of thalamic volumes across sequences. Pair-wise correlations of thalamic volumes across MPRAGE variants. Thalamic volume correlations for pairs of sequences are presented as color-coded Pearson’s correlation coefficients (r, FreeSurfer data). For each correlation matrix, sequence acquisition times (min:sec) are displayed on the axes (8:52, MP2RAGE; 6:03, meMPRAGE; 5:32, MPRAGE; 3:40, CS-MP2RAGE; 2:04 and 1:14, CS-MPRAGE). Pearson’s r interval, shown in legend, ranges from 0 to 1. Thalamic nuclei labels as in Supplementary Table 1. Pairwise correlations in FastSurfer data are presented in Supplementary Figure 3.

#### 3.3.1 Whole thalamus

In the whole thalamus, we observed excellent volumetric correlations for all pairs of sequences (Pearson’s *r* range = [0.72, 0.97]). The lowest values of *r* was observed between MPRAGE and CS-MPRAGE (TA = 2:04) data, while the highest between the two versions of CS-MPRAGE (TA = 2:04 and 1:14).

#### 3.3.2 Thalamic nuclei

In 9 out of 14 thalamic nuclei we found strong correlations regardless of sequence pair (Pearson’s *r* range = [0.45, 0.96]). Moderate correlations were found in the MD nucleus only between standard MPRAGE and CS-MP2RAGE volumes (*r* = 0.36), as well as in LGN in four pairs of sequences, all involving standard MPRAGE data. Furthermore, the weakest correlations were almost exclusively involving standard MPRAGE, mainly in smaller nuclei AV (*r* = 0.26), VM (*r* = 0.26), and MV(Re) (*r* = 0.25), as well as on the PU nucleus (*r* = 0.26).

### 3.4 Agreement between FreeSurfer and FastSurfer

In the present study, we also investigated segmentation effects according to the two segmentation tools FreeSurfer and FastSurfer. Overall, we observed good qualitative agreement between the two segmentations, despite slight differences mainly in lateroventral territories of the thalamus in the majority of participants. Figure 7 shows the results of the Bland-Altman analysis comparing thalamic nuclei volumetric data as segmented by FreeSurfer or FastSurfer. Relevant normalized volume differences were observed only in the small MGN and LD nuclei, the highest differences being on standard MPRAGE data for both nuclei. Considering all sequences together, ICC between FreeSurfer and FastSurfer thalamic volumes was 0.998 (95% CI [0.998, 0.999]). In individual sequences, it ranged between 0.996 and 1, the lower value being with CS-MP2RAGE data and the higher value with MP2RAGE.

**Figure 7:**
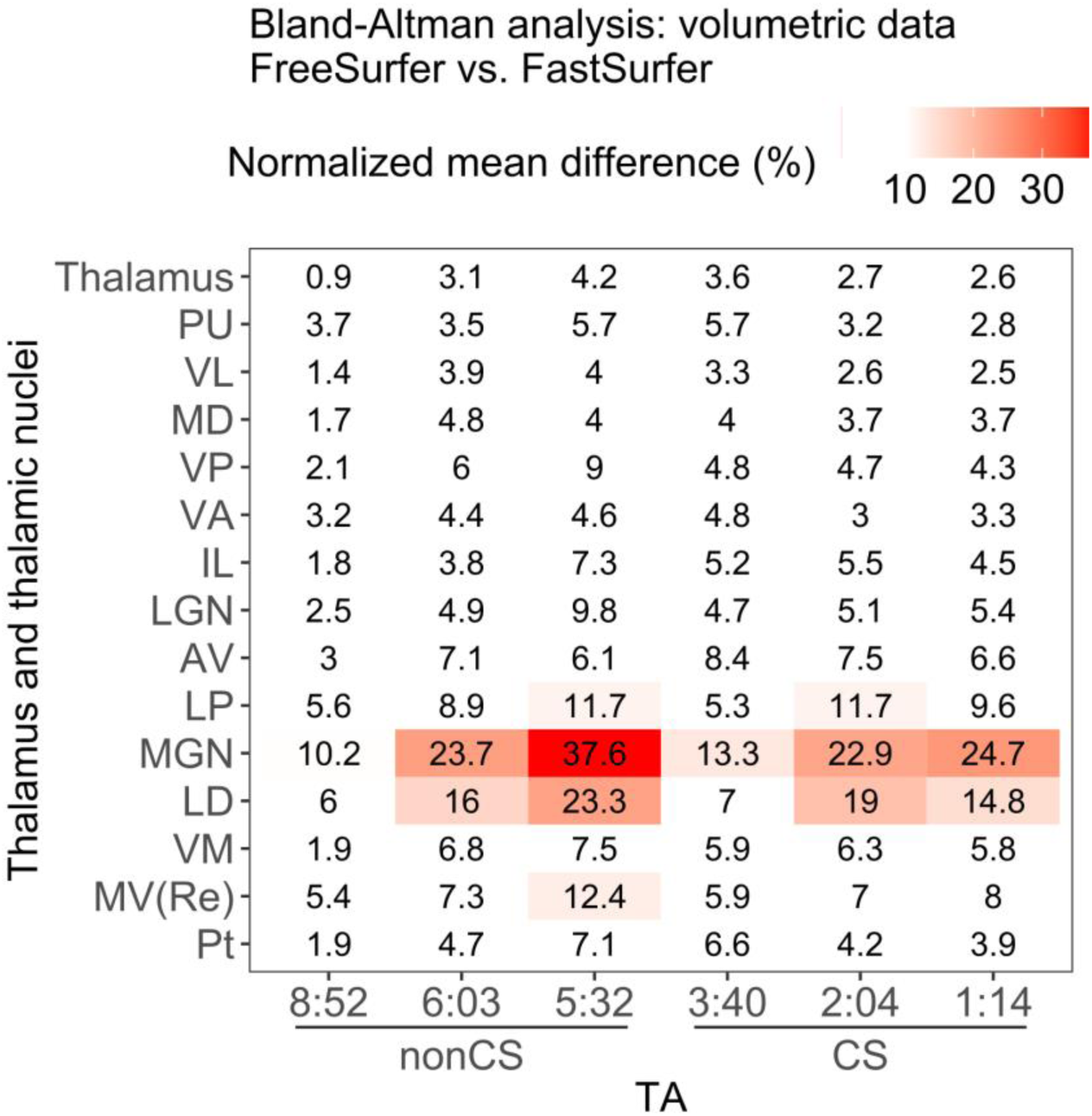
Segmentation effects, Bland-Altman analysis. Within-subject and within-sequence comparison of FreeSurfer and FastSurfer segmentation tools across MPRAGE variants and thalamic structures. Bland-Altman analysis comparing thalamic volumes from the two considered segmentation tools. For each thalamic structure, normalized mean differences were computed as the ratio of the absolute difference between volumes of the two tools to their mean, for each subject and then group averaged. Here, normalized mean differences are reported as the mean across subjects, expressed as percentage, and color-coded. Only volume variabilities higher than 10% were colored, to better highlight the highest differences (notably, nuclei MGN and LD, mean volumes across sequences of 126 and 30 mm^3^, respectively). On the x-axis, sequences are sorted in descending order of acquisition time (TA), and grouped as non compressed sensing (nonCS) and compressed sensing (CS); on the y-axis, thalamic structures are sorted in descending order of mean volume across sequences. Thalamic nuclei labels as in Supplementary Table 1.

## 4. Discussion

In the present study, we found that T1-weighted MPRAGE variants, from about 9 to 1 minute acquisition times and accelerated with the compressed sensing (CS) technique, provide human brain images with adequate quality to enable automated segmentation of subcortical thalamic nuclei at 3T. In particular, we show that, despite some degree of within-subject variation across sequences, higher accelerations are able to provide thalamic volumetric data of comparable magnitude to standard MPRAGE variants. In addition to this, we found high correlations of thalamic nuclei volumes across sequences, despite some lower correlations particularly in smaller nuclei and with data from the standard MPRAGE sequence. Furthermore, we found that the novel segmentation tool FastSurfer can be an effective, faster alternative to its counterpart FreeSurfer in the volumetric analysis of thalamic nuclei.

### 4.1 T1-weighted MRI Quality Assurance

A quality assurance of the considered MPRAGE variants revealed adequate quality of brain MR images. As expected, this result is in agreement with previous evidence covering the cortex and subcortical regions, including the whole thalamus (Mair et al., 2019, 2020; Mussard et al., 2020; Dieckmeyer et al., 2021). To the best of our knowledge, there is currently a lack of evidence concerning volume variations as a function of CS accelerations of MPRAGE sequences in thalamic nuclei specifically. Hence, the present study fills this gap showing that good image quality can be achieved with CS accelerations also in structures as relatively small as thalamic nuclei, whose borders can be particularly elusive to 3T structural MRI contrasts and parcellation techniques (Magnotta et al., 2000; Iglesias et al., 2018; Najdenovska et al., 2019; Su et al., 2019; Rushmore et al., 2022).

As far as segmentation quality and volumetry of thalamic nuclei are concerned, we found good correspondence with the literature (Benedict et al., 2013; Schoonheim et al., 2015; Iglesias et al., 2018; Shin et al., 2019). We observed some degree of variability in segmentation quality, particularly in standard MPRAGE data. It is known that the standard MPRAGE sequence performs suboptimally when separating cerebral tissues with not only T1, but also T_2_* and proton density differences (Marques et al., 2010; Mussard et al., 2020), and provides insufficient contrast to delineate thalamic nuclei specifically (Sudhyadhom et al., 2009; Tourdias et al., 2014; Tohidi et al., 2023). We hypothesize that the volumetric variability we observed on standard MPRAGE data relates to its suboptimal contrast at grey-white matter interfaces particularly, a factor that could have driven the segmentation algorithm towards a misclassification of voxels, labeling, for instance, voxels of the internal capsule as belonging to the thalamic grey matter. With respect to other contrasts, this voxel misclassification on standard MPRAGE data specifically could have produced differences mainly along lateral thalamic boundaries, overshooting voxel labels into white matter territory. This explanation is also in line with what has been stated by the developers of the thalamic parcellation tool we employed (Iglesias et al., 2018). Furthermore, these observations are consistent with the lower correlations we found in this study between standard MPRAGE data and the other sequences, for example, in voluminous nuclei such as LGN, PU, and VP, all rich in fibers or placed directly at grey-white matter interfaces (Grieve et al., 2000; Jones, 2007; Iglesias et al., 2018). As far as other MPRAGE variants are concerned, segmentations proved to be robust and of comparable quality among CS-accelerated sequences, as did standard MP2RAGE and meMPRAGE data.

Altogether, these quality assurance results suggest that the volumetry of thalamic nuclei can be investigated effectively also with CS-accelerated MPRAGE variants at acquisition times of about 1-2 minutes, as valid substitutes to longer and more conventional MPRAGE sequences.

### 4.2 Within-Subject Sequence Effects

We found that the choice of MPRAGE variants introduces variability in the volumes of thalamic nuclei. More specifically, coefficients of variation (CV; FreeSurfer-data) were above 10% in a third of the considered thalamic structures, consisting of smaller nuclei.

It is known that smaller brain structures, such as hippocampus and amygdala, which like the thalamus have a predominant composition of grey-white matter interfaces, exhibit high volume variabilities in volumetric studies (Bartzokis et al., 1993; Pruessner et al., 2000; Mueller et al., 2007), particularly when MR images differ in acquisition methods (Seiger et al., 2021). We hypothesize that the volume variations we observed in thalamic nuclei are related to their size, rather than being driven by CS accelerations. Indeed, the fact that we observed volume variations of less than 10% in bigger nuclei and less than 5% in the whole thalamus suggests that variations could relate to the size of nuclei, a result further supported by the negative correlation we found between CV and mean volumes of thalamic nuclei.

We also performed a volumetric analysis considering non CS and CS data separately. We observed lower volume variations with CS data generally, notably in smaller thalamic nuclei as well. We also found a higher proportion of significant sequence effects in non CS data as compared to CS data. Altogether, these results suggest higher volume variabilities particularly in non CS data and small thalamic nuclei. We speculate that these results are due to the different MPRAGE variants used in this study. The non CS sequences, namely MPRAGE, meMPRAGE, and MP2RAGE, are quite heterogeneous in terms of acquisition methods: MPRAGE is composed of only one single-echo T1-weighted image (Mugler & Brookeman, 1990), meMPRAGE results from averaging four different echoes (Van der Kouwe et al., 2008), and the MP2RAGE uniform image is produced with a particular combination of volumes that rules out B_1_ contributions to the image, using a single-echo approach (Marques et al., 2010). On the other hand, a comparison of thalamic nuclei volumetry across the CS sequences in this study, namely CS-MP2RAGE and the two versions of CS-MPRAGE, are all single-echo and differ mostly in the *k*-space undersampling factor which shortens acquisition times. Even if some degree of variability in the acquisition methods is preserved across the CS group as well, since the CS-MP2RAGE acquisition method differs from CS-MPRAGE, the fact that two out of three sequences in the CS group are almost identical could have driven the lower volume variability that we observed here, in contrast to the non CS group, more heterogeneous in terms of acquisition protocols.

In addition to this, we found significant sequence effects for all the considered thalamic nuclei. However significant the volume variability across MPRAGE variants may be in our study, the effect sizes of sequence effects were all small (Kendall’s W ≤ 0.2). This result indicates that sequence effects might play a little role in the observed volume variability altogether. In fact, it further suggests that the volume variability could ensue from a combination of acquisition methods, particularly in relation to non CS sequences, and anatomical characteristics of thalamic nuclei, their size and white matter fraction being the most prominent factors.

In the light of this, the volume variability in thalamic nuclei that we observed could potentially be attributed to the combined effects of nuclei sizes and choice of CS acceleration. Nonetheless, the fact that the whole thalamus and major thalamic nuclei were less variable than minor nuclei, and that the observed sequence effects were little meaningful, suggests that the use of CS-accelerated MPRAGE sequences can be an effective approach in the volumetric characterization of thalamic nuclei.

### 4.3 Volume Correlations

We also evaluated volume correlations across pairs of MPRAGE variants in the considered thalamic nuclei (FreeSurfer data). The strongest correlations were seen in the whole thalamus, major nuclei VL, VP, VA, and IL, as well as in minor nuclei LP, MGN, LD, and Pt across all pairs of sequences. It is interesting to note that weak correlations were observed in VP nucleus when standard MPRAGE data was considered, a result further suggesting that this sequence might not be the first-choice approach when characterizing thalamic nuclei structurally with MRI. The weakest correlations for pairs of MPRAGE variants were on minor nuclei AV, VM, and MV(Re) in different combinations of sequences, as well as on major nuclei LGN, MD and PU, again notably involving standard MPRAGE data in these three latter cases.

As previously mentioned, standard MPRAGE is known to perform suboptimally when separating tissues with different properties in certain brain regions (Marques et al., 2010; Mussard et al., 2020). Our correlational results, specifically in nuclei VP, PU, MD, and LGN when standard MPRAGE data was involved, suggest that the lower contrast that this sequence provides, for instance, in portions of the thalamus interfacing white matter structures (e.g. internal capsule, thalamic radiations, and internal medullary lamina), could bias the segmentation of these nuclei, hence subsequent volume correlations. Alonso et al. (2021) and Ferraro et al. (2022) found evidence that the volume of the whole thalamus can change significantly in relation to the choice of MPRAGE variants. These authors included standard MPRAGE in their comparisons with a single other CS-accelerated MPRAGE variant, and did not investigate sequence effects in thalamic nuclei exclusively, by reason of their main interest in other cortical and subcortical structures. In the light of all this, the choice of standard MPRAGE sequence as the only comparison term with CS-accelerated ancillary sequences might not be optimal for analyses in the thalamus area. Our data suggest that other sequence variants could be considered or included to fully appreciate potential differences across sequences, especially in studies including thalamus and/or thalamic nuclei.

The highest correlation on nearly all the considered thalamic nuclei was between the two CS-accelerated MPRAGE sequences. As previously mentioned, they have an almost identical acquisition method, their only difference being *k*-space sampling factor. This characteristic could explain the high correlation values across thalamic nuclei observed between these two CS-accelerated MPRAGE sequences.

The low correlations we observed in the PU nucleus were a puzzling result. Since it is one of the most voluminous nuclei in the primate thalamus (Grieve et al., 2000; Ferris et al., 2013), we expected its volume to be less variable across sequences. On the contrary, it yielded some of the lowest correlations of the dataset. In this regard, we hypothesize that the grouping procedure we used to merge the PU subdivisions into a single entity could be at the basis of the observed lower correlations in this nucleus specifically. By performing separate analyses of the four subdivisions of PU, we found that the subdivisions PuM, PuA, and PuI, but not PuL, had weak correlations involving standard MPRAGE data, with correlations going slightly below zero in the subdivision PuA between standard MPRAGE and MP2RAGE, and standard MPRAGE and CS-MP2RAGE (data not shown). This finding could relate partly to the aforementioned limitation of standard MPRAGE when segmenting different tissues, especially when compared, for instance, to the excellent contrast provided by MP2RAGE sequences. Furthermore, given the high range of cortico-pulvino-cortical connections subserving a wide variety of cognitive functions (Sherman & Guillery, 2006; Fiebelkorn & Kastner, 2020), we speculate that a differential proportion of white matter fibers could innervate the four subdivisions of the PU nucleus. By virtue of the poorer contrast of standard MPRAGE data, the differential proportion of white matter fibers in nucleus PU could ultimately have yielded weaker correlations when standard MPRAGE data, again, was involved.

### 4.4 Agreement between FreeSurfer and FastSurfer

To the best of our knowledge, we provide first evidence that thalamic parcellations (Iglesias et al., 2018) can be obtained from FastSurfer-segmented (Henschel et al., 2020) data as well, in addition to segmentations from the FreeSurfer software, as mentioned by the authors of the thalamic nuclei parcellation tool in their guidelines (Iglesias et al., 2018). We performed a comparison between volumetric data of the considered thalamic nuclei as segmented by FreeSurfer or FastSurfer, and observed overall good qualitative agreement between the two tools. Bland-Altman analysis revealed notable differences only in the small MGN and LD nuclei, the highest differences being on standard MPRAGE data, while excellent ICC suggests a very high consistency between the two segmentation tools for all sequences. FastSurfer has been recently proposed as an alternative to the intensive-runtime FreeSurfer segmentation tool, providing comparable volumetric results to its counterpart in a number of brain regions (Henschel et al., 2020; Bloch & Friedrich, 2021; Kemenczky et al., 2022; Müller et al., 2023), including the whole thalamus (Opfer et al., 2023). Our findings are in line with the literature and extend their validity to the thalamic nuclei specifically.

Taken together, our findings suggest that the characterization of thalamic nuclei with 3T MRI in humans could benefit from both the use of CS accelerations and the segmentation tool FastSurfer, whose combined effect can optimize the schedules required in multiple steps of MRI volumetric studies in terms of time consumption.

Furthermore, by showing the feasibility of structural and volumetric analysis with MRI data taken at short acquisition times, our findings pave the way to future studies aiming to characterize thalamic nuclei structural, diffusional, connectional, and functional profiles in either healthy subjects or in highly kinetic populations, such as, for instance, elderly people and patients (Van Dijk et al., 2012; Iglesias et al., 2017; Madan, 2018; Noor et al., 2020).

## 5. Conclusions

We show the feasibility of automatic thalamic nuclei segmentation with 3T T1-weighted MPRAGE variants using different degrees of compressed sensing (CS) acceleration in a sample of healthy adults.

Although within-subject thalamic volumes are affected by the choice of sequences, volume variability is low for the whole thalamus and major nuclei, and volume correlations are of appreciable magnitude for the majority of the considered structures. Additionally, the choice of the segmentation tool FastSurfer could represent a robust alternative to its counterpart FreeSurfer in order to further reduce times in the segmentation of the thalamic nuclei. Our study also shows that, although the effect sizes can be small, the choice of MPRAGE sequence variant matters because it affects the segmentation of thalamic nuclei. This means that in retrospective or prospective MRI studies, attention should be placed to minimizing sequence differences (both within and across MRI sites), or to include these effects as part of the analyses.

Based on our evidence, future studies can employ CS-accelerated MPRAGE variants to characterize thalamic nuclei, along with multimodal imaging methods and in different populations, particularly those in which shorter acquisition times are required in order to reduce the possibility of distortions introduced by high level of motion.

## Data and code availability statement

Data are available via request to the authors, with the need of a formal data sharing agreement.

## Acknowledgments

This work was supported by funding from the Municipality of the City of Rovereto (Trento), Italy, for the project *“Advanced neuroimaging to study aging”*.

---

**Supplementary Figure 1.**
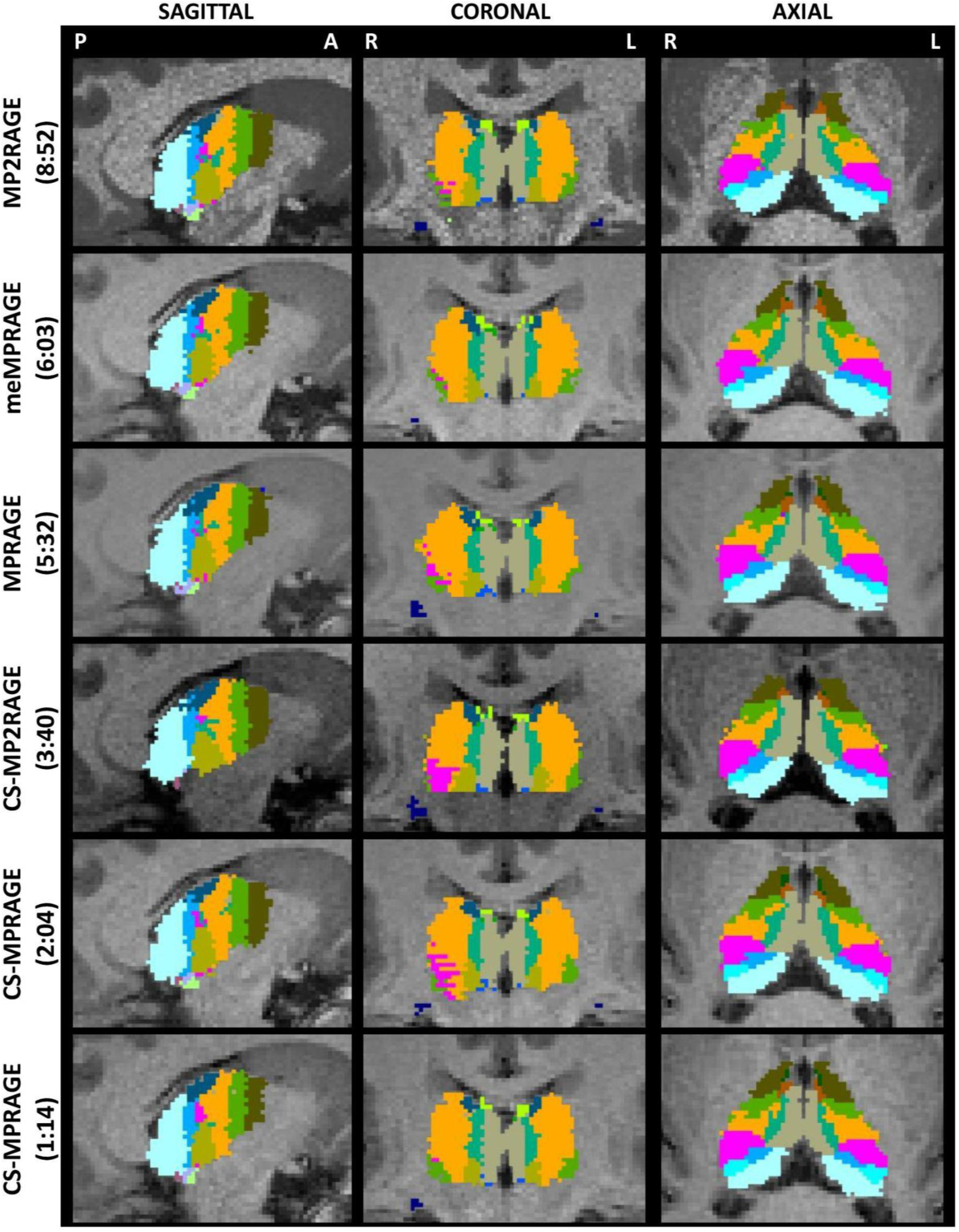
FastSurfer data: Thalamic nuclei segmentations for differently accelerated MPRAGE variants. FastSurfer automated thalamic nuclei parcellations of a representative subject across the considered MPRAGE variants (acquisition times in brackets, sequences details in Table 1). Sagittal, coronal, and axial planes of parcellations are with the same brightness parameters as in Figures 1 and 3. As in those figures, slices are taken at central thalamus (subject’s native space). Sagittal images show the right hemisphere. Nuclei labels and colors as in Figure 2 and Supplementary Table 1. FreeSurfer segmentations on the same subject’ can be seen in Figure 3.

**Supplementary Figure 2.**
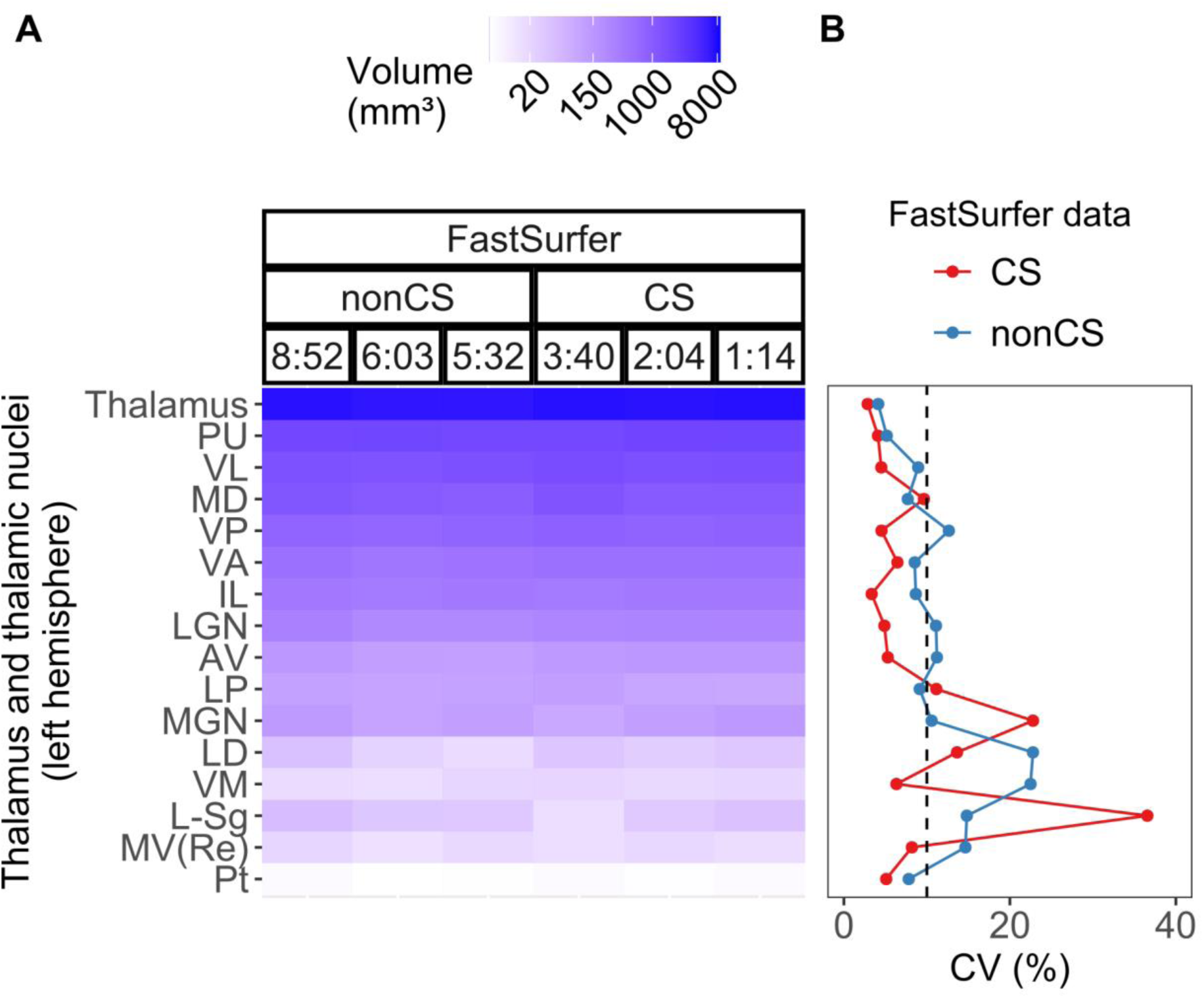
FastSurfer data: Within-subject volume variation in thalamic structures across MPRAGE variants. A) Thalamic volumes (logarithmic color-coding by size) segmented by FastSurfer (left hemisphere) for each MPRAGE variant, listed by decreasing acquisition times (TA, minutes:seconds) and grouped as non compressed sensing (nonCS) and compressed sensing (CS). B) Group average of within-subject coefficients of variation (CV) across MPRAGE sequences for thalamic structures (left and right hemispheres averaged), considering CVs from nonCS (blue) and CS (red) sequences separately. Sequence TA (minutes:seconds): 8:52, MP2RAGE; 6:03, meMPRAGE; 5:32, MPRAGE; 3:40, CS-MP2RAGE; 2:04 and 1:14, CS-MPRAGE. The vertical dash line corresponds to a reference CV of 10%.

**Supplementary Figure 3.**
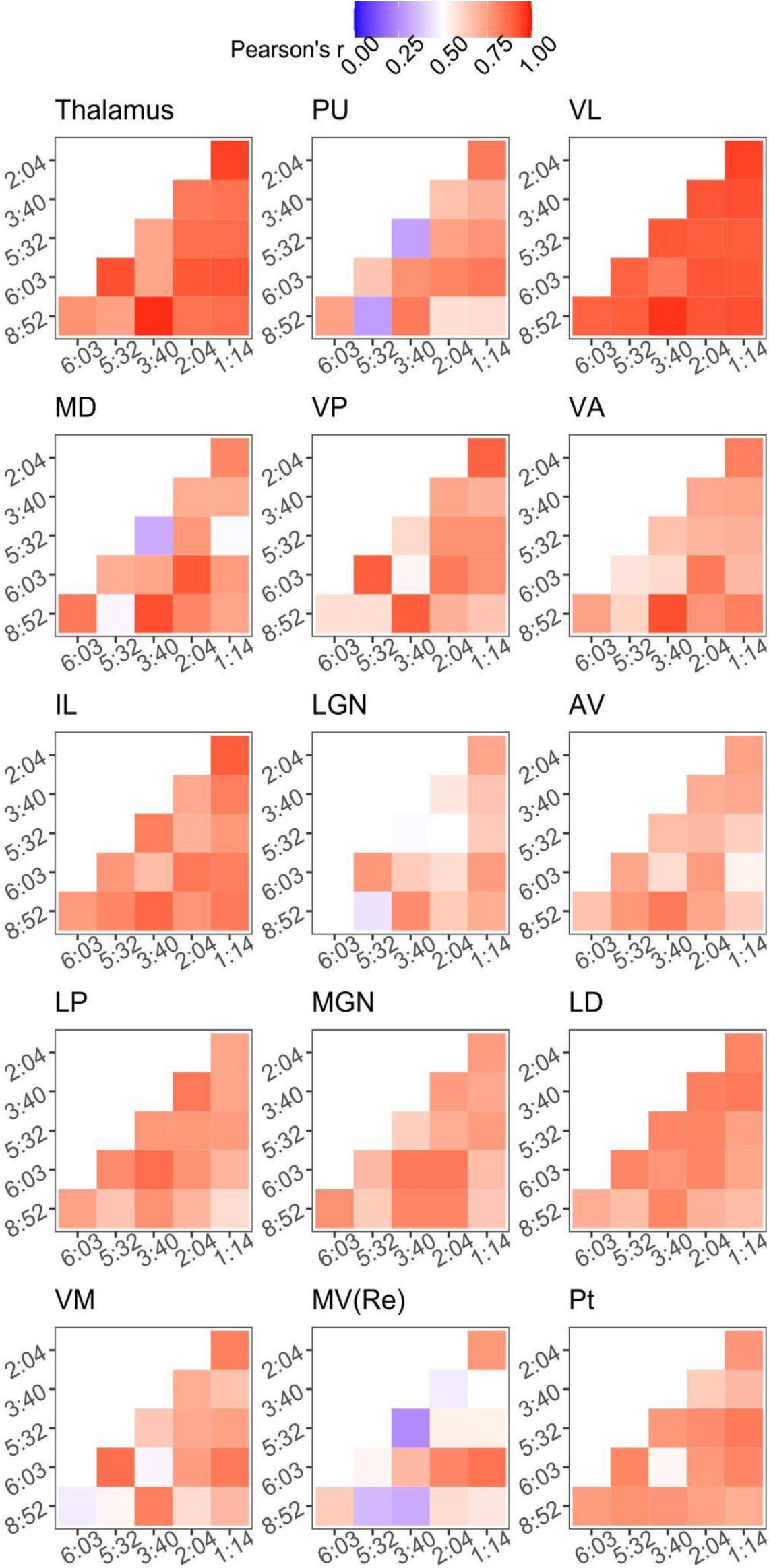
Pair-wise correlations of thalamic volumes across MPRAGE variants, FastSurfer data. Thalamic volume correlations for pairs of sequences are presented as color-coded Pearson’s correlation coefficients (r). For each correlation matrix, sequence acquisition times (min:sec) are displayed on the axes (8:52, MP2RAGE; 6:03, meMPRAGE; 5:32, MPRAGE; 3:40, CS-MP2RAGE; 2:04 and 1:14, CS-MPRAGE). Pearson’s r interval, shown in legend, ranges from 0 to 1. Thalamic nuclei labels as in Supplementary Table 1. Pair-wise correlations in FreeSurfer data are presented in Figure 6.

**Supplementary Table 1.**
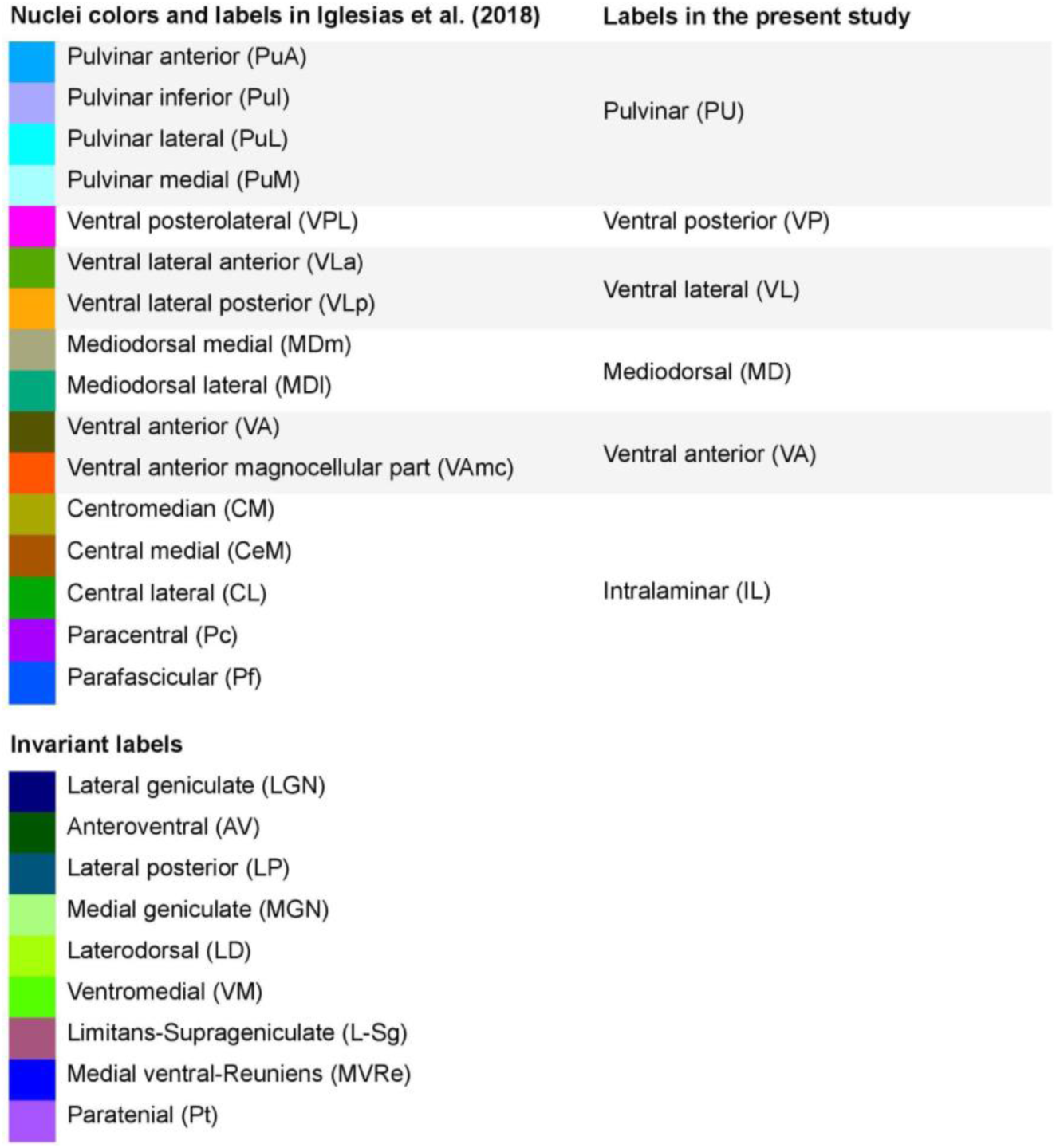
Thalamic nuclei colors, original labels (Iglesias et al., 2018), and re-labeling scheme used in the present study. Similar re-labeling can be found in Iglesias et al. (2018), Bocchetta et al. (2020), Tregidgo et al. (2023).

| Nuclei colors and labels in Iglesias et al. (2018) |  | Labels in the present study |
| --- | --- | --- |
|  | Pulvinar anterior (PuA) | Pulvinar (PU) |
|  | Pulvinar inferior (Pul) |  |
|  | Pulvinar lateral (PuL) |  |
|  | Pulvinar medial (PuM) |  |
|  | Ventral posterolateral (VPL) | Ventral posterior (VP) |
|  | Ventral lateral anterior (VL<sub>a</sub>) | Ventral lateral (VL) |
|  | Ventral lateral posterior (VL<sub>p</sub>) |  |
|  | Mediodorsal medial (MD<sub>m</sub>) | Mediodorsal (MD) |
|  | Mediodorsal lateral (MD<sub>l</sub>) |  |
|  | Ventral anterior (VA) | Ventral anterior (VA) |
|  | Ventral anterior magnocellular part (VA<sub>mc</sub>) |  |
|  | Centromedian (CM) | Intralaminar (IL) |
|  | Central medial (CeM) |  |
|  | Central lateral (CL) |  |
|  | Paracentral (Pc) |  |
|  | Parafascicular (Pf) |  |
| **Invariant labels** |  |  |
|  | Lateral geniculate (LGN) |  |
|  | Anteroventral (AV) |  |
|  | Lateral posterior (LP) |  |
|  | Medial geniculate (MGN) |  |
|  | Laterodorsal (LD) |  |
|  | Ventromedial (VM) |  |
|  | Limitans-Suprageniculate (L-Sg) |  |
|  | Medial ventral-Reuniens (MVR<sub>e</sub>) |  |
|  | Paratenial (Pt) |  |

